# Neural Predictors of Functional Genomic Responses to Negative Social Evaluation in Adolescent Females

**DOI:** 10.1101/2025.06.18.660364

**Authors:** Mathilda von Guttenberg, Jeffrey Gassen, George M. Slavich

## Abstract

**Background:** Social stress—particularly when experienced during adolescence, can have a lasting impact on health and well-being. Among other key biological pathways, inflammatory and innate immune signaling appear to play important roles in linking stress to physical and mental health problems. Individual differences in sensitivity to social threats may leave certain people more vulnerable to stress and its harmful sequelae than others, and a growing body of research has found that stress sensitivity is reflected in neural activity throughout the threat network. However, few studies have investigated whether heightened neural sensitivity to social threats is related to acute changes in immune and neuroendocrine pathways relevant to health, particularly among those for whom the effects of stress may be especially impactful.

**Method:** In the current research, 52 adolescent females (*M_Age_ =* 14.90*, SD =* 1.35) participated in a functional magnetic resonance imaging study to examine brain activity and functional connectivity during a social evaluation task. Nearly half of the sample (*n* = 22) were identified as having a maternal history of depression. Blood samples were collected prior to the task, as well as 35 and 65 min. after the task began, and were used for transcriptional profiling.

**Results:** The primary analyses tested whether threat network activity and connectivity predicted the magnitude of change in gene expression from baseline to the follow-up time points. Results revealed robust shifts in expression of genes in innate immune pathways in response to the task (e.g., hypoxia inducible factor-1, interferon signaling). Although activity across the entire threat network was related to individual differences in gene expression, anterior cingulate cortex-insula and insula-ventromedial prefrontal cortex connectivity were most consistently related to up- and down-regulation of immune pathways, respectively. These patterns were further moderated by differences in maternal depression history.

**Conclusion:** Results demonstrate that individual differences in threat network activity may have important implications for biological responses to social threat among adolescent females. In turn, these findings both provide insights into neural signatures of social stress vulnerability and the biological pathways that may contribute to poorer health outcome among those most vulnerable to stress.

## 1. Introduction

Social stress has a profound impact on mental health, ^1–3^ with experiences of social rejection emerging as one of the strongest proximal predictors of major depressive disorder (MDD) across the life course. ^4–6^ Adolescence is a particularly vulnerable period for these effects, as peer relationships and social hierarchy become increasingly important during this time period. ^7^ Prior to the pubertal transition, MDD rates are similarly low for males and females; following the pubertal transition, however, rates of MDD approximately double for females, with the reasons for these effects still being unclear. ^8, 9^ Consequently, additional research is needed to understand factors that may underlie the striking increase in MDD risk for adolescent females, especially if such research can help identify *potentially modifiable preclinical risk factors* for the disorder.

According to the Social Signal Transduction Theory of Depression, social stress may have lasting effects on both mental and physical health by heightening neural sensitivity to social threat and by activating innate immune processes involved in inflammation, ^10, 11^ which in turn impacts health. ^12, 13^ In vulnerable individuals, this biological response may contribute to the onset of depressive symptoms and increase the risk of developing comorbid somatic conditions frequently observed alongside MDD. ^11, 14–16^ However, the precise mechanisms linking social rejection with risk for depression and related conditions remain unclear.

### 1.1 Neural risk processes in depression

From a neurobiological perspective, adolescent social stress may influence depression risk by altering the activity and/or connectivity of brain networks involved in stress reactivity and inflammation regulation. ^17–19^ As posited by the Social Signal Transduction Theory of Depression, ^11^ experiences of social rejection are converted into neural signals of threat in a set of brain regions, commonly referred to as the threat network, which includes the amygdala, anterior insula, and anterior cingulate cortex (ACC)—areas highly responsive to both physical and social threats (for review, see ref. ^11^). These changes may be further reflected in altered functional connectivity between the threat network and regions involved in negative affect and emotion regulation, such as the ventromedial prefrontal cortex (VMPFC). ^20, 21^

An implication of this framework is that individuals at increased risk for depression—for instance, those with a maternal history of major depressive disorder—may exhibit stronger neural responses to social rejection than those without this predisposing risk factor.^2^ Supporting this notion, past research has shown that heightened activity in threat network regions is observed in depression,^22, 23^ correlates with greater depression severity, ^24, 25^ and is associated with heightened neuroendocrine reactivity to stress,^26^ and inflammatory responding.^27^ Together, these findings suggest that the threat network may serve as a key pathway linking social stress and risk for depression in adolescence. However, for neural activity to drive widespread physiological changes, it must trigger a biological cascade that extends beyond the brain, embedding social stress in the periphery and shaping immune and inflammatory pathways that contribute to depression risk.

While much of the existing literature has focused on the threat network as a key neural pathway linking social stress to depression risk, recent meta-analyses exploring the neural underpinnings of social stress processing in humans ^28^ have identified additional brain regions such as the thalamus, claustrum, inferior frontal gyrus, and posterior cingulate cortex. ^29–31^ These findings may reflect the diversity of experimental paradigms used to elicit social stress, as brain activity tends to be highly task-dependent. Given this methodological variability across studies, a brain-wide mapping approach is preferable—not only to avoid prematurely constraining analyses to predefined regions of interest, but also to identify novel or unexpected neural contributors to social stress processing.

### 1.2 Inflammation and depression

In terms of peripheral biology, growing evidence indicates that pro-inflammatory signaling pathways may play a key role in linking social stress to depression risk.^32^ Social rejection has been shown to activate the innate immune and, in particular, inflammatory response, resulting in elevated levels of pro-inflammatory cytokines—biomarkers frequently observed to be elevated in individuals with depression.^11, 33^ These heightened cytokine levels are not only associated with depression but also with several physical health conditions that commonly co-occur with MDD, including asthma, heart disease, chronic pain, and autoimmune and neurodegenerative disorders.^12, 34^

At the genomic level, social stressors, including social rejection, have been associated with upregulation of pro-inflammatory immune response genes including *IL1B, IL6, IL8, TNF*, and downregulation of antiviral immune response genes including *IFNA and IFNB*, a pattern that has been referred to as the Conserved Transcriptional Response to Adversity.^10, 11, 32, 35^ This transcriptional shift is thought to be mediated by increased activity of pro-inflammatory transcription factors such as NF-κB and AP-1, along with reduced glucocorticoid receptor sensitivity, which acts to regulate inflammation among other functions critical to health.^32^

The central nervous system (CNS) appears to play a pivotal role in translating social threat signals into biological changes through social signal transduction pathways.^11^ For example, the experience of social stress or rejection activates the sympathetic nervous system (SNS) and hypothalamic-pituitary-adrenal (HPA) axis, leading to the release of neurotransmitters and hormones, such as epinephrine/norepinephrine and cortisol.^36, 37^ In turn, these chemical messengers interact with receptors on immune cells to induce transcriptional changes that impact inflammatory signaling, cell proliferation and maturation, and core effector functions.^11, 38, 39^

Importantly, not all individuals exhibit the same biological sensitivity to stress, with resulting implications for their risk of developing MDD.^40, 41^ In evidence of this, recent research using an endotoxin challenge in healthy participants found that individual differences in perceived stress, sensitivity to social disconnection, and preexisting depressive and anxiety symptoms moderated the effects of endotoxin exposure on both mood and pro-inflammatory gene expression.^42^ We previously published the first study linking neural responses to social rejection with inflammatory responses to social stress in young adults.^27^ What remains unknown, however, is whether individual differences in neural sensitivity to social rejection influence biological responses to social stress in adolescence, potentially affecting subsequent depression outcomes in this high-risk developmental period. Understanding how variation in neural responsivity contributes to the stress-immune response at the transcriptional level, in particular, could clarify why some youth are more vulnerable to inflammation-related mood disturbances, thus providing new insights into the pathophysiology of MDD.

### 1.3 Present Study

As described by Slavich and Irwin (2014),^11^ substantial evidence supports the core principles of the Social Signal Transduction Theory of Depression (see also ^16, 43–46)^. However, neural and genomic processes to acute social stress have very rarely been examined together in the same individuals, with most of the existing research relying on separate lines of investigation, thus leaving key questions about their interrelation unresolved. Moreover, research on acute transcriptional changes following a social stressor task remains limited, particularly in adolescent females at varying risk for depression. Finally, to our knowledge, few studies to date have examined whether social evaluation-induced neural activity and connectivity in threat-related brain regions predict transcriptional changes in immune and inflammatory pathways, making this a critical gap in understanding the biological pathways linking social stress to risk for MDD in youth.

To address gaps in our understanding of risk for depression in adolescence, we conducted the Psychobiology of Stress and Adolescent Depression (PSY SAD) Study. ^2^ In the present study, we extend work on this well-characterized cohort by investigating the extent to which activity and functional connectivity in key threat-related brain regions—namely, the amygdala, anterior insula, ACC, VMPFC—predict transcriptional changes to an fMRI-based acute social stressor. Second, we examined whether neural-genomic associations were moderated by youths’ risk for depression by comparing the effects of neural sensitivity on transcriptional changes to the social stressor for adolescent females at high vs. low risk for developing MDD based on their mothers’ lifetime history of the disorder. Finally, we conducted a whole-brain voxel-wise analysis to explore whether stress-related transcriptional responses were associated with broader patterns of neural activity across the brain.

Based on the Social Signal Transduction Theory of Depression,^11^ we had two primary hypotheses. First, we hypothesized that neural responses to negative social evaluation, specifically activity and connectivity within the amygdala, insula, ACC, and VMPFC, would predict transcriptional changes in immune and inflammatory pathways, with distinct patterns of connectivity (e.g., ACC–insula vs. insula–VMPFC) associated with up- or down-regulation of gene expression.^27^ Second, we hypothesized that these neural–genomic associations would differ by depression risk status, with high-risk adolescents showing stronger or altered transcriptional responses relative to low-risk peers.

## 2. Methods [See Ref. ^2^]

### 2.1 Participants

Participants in the PSY-SAD study^2^ (*N =*52*, M_Age_ =* 14.90*, SD =* 1.35) were recruited using online advertisements, flyers posted in community locations, social media posts, word of mouth and announcements made at middle and high schools in the greater Los Angeles area.^2^ To qualify for the study, at the time of recruitment, daughters had to be between the ages of 12 and 16, English-speaking, right-handed, not claustrophobic, free of bodily metal (except dental fillings) and other contraindications for MRI, living with their biological mother and have no current or past history of any Diagnostic and Statistical Manual-IV (DSM-IV) Axis I disorder. In addition, daughters were required to meet the following criteria: no recent history of alcohol or substance use or dependence, no confirmed pregnancy as verified by a pregnancy test, and no history of head trauma or learning disabilities. Furthermore, participants needed to be free from factors known to affect inflammation, such as past or current inflammatory illnesses, significant sleep disturbances, tobacco use, prescription medication use, excessive caffeine consumption (defined as more than eight beverages per day), or a body mass index (BMI) of 30 or higher.^47^

### 2.2 Recruitment and Intake

Interested mothers and daughters participated in a phone screening followed by an in-person intake session if deemed eligible. During the intake session, written informed consent was obtained from mothers, while daughters provided assent. Both mothers and daughters were separately screened by trained diagnostic interviewers, who were all trained by George M. Slavich, to ensure they met the inclusion/exclusion criteria, and to classify the daughters’ maternal risk group. A history of maternal depression is linked to both an earlier onset and greater severity of depression in daughters, making it the determinant of risk classification in this study.^48^ Maternal risk was categorized as either low-risk (mothers with no history of major depressive episodes [MDEs]; N = 30) or high-risk (mothers with at least one MDE; N = 22).^2^ Once the diagnostic interviews were completed, daughters completed self-report questionnaires assessing their demographics (see **Table 1**). Shortly after, daughters completed a 10-minute video-recorded impressions interview, adapted from Muscatell et al. (2015), in which they discussed their upbringing, opinions, feelings, and autobiographical memories while alone (i.e., without their mothers present).^49^ The interview consisted of 33 open-ended, conversationally framed questions, designed to elicit personally meaningful responses. These included items such as: “What is your favorite hobby?”, “What are you most afraid of?”, and “What qualities do you value most in a friendship?”. Additional procedural details are provided in Supplemental Table 1 and in Ref.^2^

**Table 1.** Sample Characteristics by MDD risk Group.

|  | High-risk female<br>adolescents<br>*n* = 22<br>mean (*SD*) | Low-risk female<br>adolescents<br>*n* = 30<br>mean (*SD*) | Group<br>difference<br>*p* |
| --- | --- | --- | --- |
| Age | 14.68 (1.39) | 15.07 (1.31) | .31 |
| Body mass index | 24.87 (6.70) | 22.24 (4.51) | .10 |
| Race |  |  | .45 |
| Black or African<br>American | 9 % | 7 % |  |
| Asian or Asian<br>American | 0 % | 7 % |  |
| White/Caucasian | 23 % | 40 % |  |
| Hispanic/Latina | 32 % | 27 % |  |
| Other | 5 % | 7 % |  |
| Mixed/multiple | 32 % | 13 % |  |

### 2.3 Study Visit Two (Experimental Session)

Following the completion of all procedures during the intake session, daughters and their mothers were scheduled for a second, three-hour experimental session. This session typically occurred within one month of the intake session (median = 26.5 days).

Upon arriving for this session, daughters were taken to private testing rooms, where they were informed of the sessions procedures and prepped for a blood draw. For the blood draws, a nurse from the UCLA Clinical and Translational Research Center (CTRC) inserted an MRI-safe indwelling catheter into the participant’s non-dominant forearm. Participants were given 15 minutes to acclimate to the catheter, after which their baseline blood sample was drawn.^2^ Soon after, daughters were introduced to “another participant,” who they were told was taking part in a related study. In reality, this “participant” was a confederate—a female, college-aged research assistant—who dressed and acted like an adolescent to create a sense of social evaluative threat for the participant.

Approximately 1 hour after the start of the experimental session, both the participant and the confederate were taken to the MRI scanner control room. Here, the participant and confederate were introduced to the Social Evaluation Task.^50^ First, participants were reminded of the recorded interview from the first session and told that the “other participant” (the confederate) would be watching their video and rating it by selecting one of 24 adjectives (divided evenly into positive, neutral, and negative categories) to describe their impressions of the participant (Fig. 1). To make the setup more convincing, the confederate asked questions such as, “How often should I provide a rating?” during the instructions. Participants were told they would see the adjective ratings in real time, but in reality, all participants viewed the same pre-recorded video showing adjectives being “selected” in a pseudorandom order. The sequence included an interval of about 10 seconds between words, with no more than two similarly valenced adjectives appearing consecutively. The task lasted 10 minutes. Neural activity and connectivity readout were time locked to adjective selection.

**Fig. 1.**
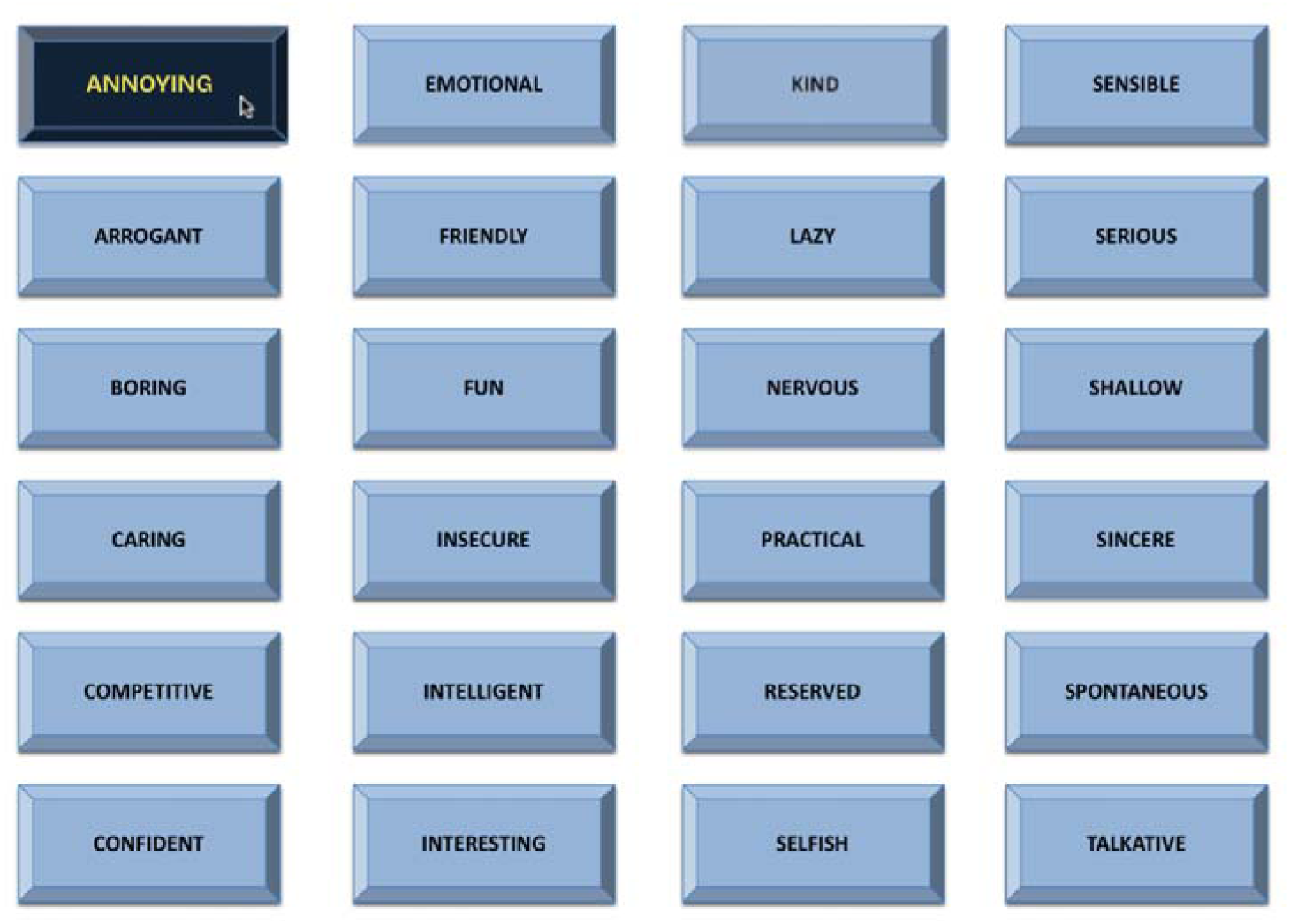
A screenshot of the social evaluation task that participants saw while in the Functional magnetic resonance imaging (fMRI) scanner.

Participants also provided post-scan blood samples at 35 and 65 minutes after the start of the Social Evaluation Task. After completing the scan, participants returned to the testing room, where they were fully debriefed on the research process and intentions of the study.

### 2.4 Blood collection & transcriptomic reprocessing

Blood was drawn once prior to the Social Evaluation Task (i.e., a baseline blood draw approximately 55 minutes before the start of the Social Evaluation Task) and twice following the task (i.e., at approximately 35 and 65 minutes after the Social Evaluation Task began). At each time point, 2.5 mL of blood was drawn into a PAXgene Blood RNA Tube to test for social stress-induced changes in gene expression using genome-wide transcriptional profiling. By the end of each study session, samples from all 3 time points were transferred to the UCLA Center for Pathology Research Services, where blood tubes were frozen at 80 C. Samples were then transferred to the UCLA Social Genomics Core Laboratory where total RNA was extracted (RNeasy;Qiagen, Valencia, CA), tested for suitable mass (Nanodrop ND1000) and integrity (Agilent Bioanalyzer), converted to barcoded cDNA (LexogenQuantSeq 30 FWD), and sequenced on an Illumina HiSeq4000 system (Illumina, San Diego, CA) in the UCLA Neuroscience Genomics Core Laboratory, all following the manufacturer’s standard protocols for this workflow. Assays targeted >10 million 65-nt single-stranded sequence reads for each sample, each of which was mapped to the reference human transcriptome using the STAR aligner and quantified as gene transcripts per million mapped reads. Analyte values were log_2_-transformed prior to analysis. (See Ref.^2^)

### 2.5 fMRI image acquisition

Imaging data were acquired using a Prisma 3.0 T whole-body scanner (Siemens Medical Systems, Iselin, New Jersey) at the Staglin One Mind Center for Cognitive Neuroscience at UCLA. High resolution T1-weighted structural images were acquired using a magnetized prepared rapid acquisition gradient echo (MPRAGE) sequence containing 1.1 mm isotropic voxels, TR/TE/flip angle =2300 ms/2.95 ms/9◦, FOV=270 mm2, 176 slices. Blood oxygenation level-dependent (BOLD) functional images were acquired containing 3 mm isotropic voxels, TR/ TE/flip angle =2000 ms/34 ms/76◦, FOV =208 mm2, 48 slices.^2^

### 2.6 fMRI preprocessing and analysis

Neural activity and connectivity estimates used in this study were derived from preprocessing and analysis pipelines detailed in Shields et al. (2024).^45^

#### 2.6.1 Activity

Functional and structural MRI data were preprocessed using SPM12 following standard pipelines described in Shields et al. (2024).^45^ Preprocessing steps included realignment and unwarping of functional images, slice-timing correction, motion correction, co-registration, segmentation, normalization, and centering. Regions of interest (ROIs) for the amygdala, anterior insula, ACC, and VMPFC were defined using a combination of the Harvard-Oxford subcortical atlas, the CONN Toolbox salience network ROIs, and a VMPFC mask from Bhanji et al. (2019).^51^ ROI-based signal intensities were extracted using REX and standardized prior to statistical analysis.

#### 2.6.2 Functional Connectivity

Functional connectivity analyses were conducted using the CONN Toolbox v19.b, following the preprocessing and denoising pipeline described by Shields et al. (2024).^45^ Preprocessing included motion correction, outlier detection via ART, segmentation, normalization, and denoising using aCompCor. Generalized psychophysiological interaction (gPPI) analyses were used to compute condition-specific functional connectivity between a priori ROIs, with contrasts reflecting differential connectivity during viewing of negative vs. neutral adjectives. Motion parameters, ART outlier flags, and physiological noise regressors were included as covariates at the first level. ROI-to-ROI connectivity measures used in this study correspond to the contrasts described in Shields et al. (2024).^45^

### 2.7 Data Analysis

To examine whether changes in gene expression following the social evaluation task were related to neural activity or connectivity across threat-related brain regions, we conducted a series of longitudinal analyses of covariance (ANCOVAs) using R software (*lme4* R package). These models estimated the extent to which change in expression of each gene (log_2_-transformed) from baseline to the follow-up time points varied depending on activity in each brain region (z-scored) during the task, controlling for baseline expression. Next, we conducted analyses a second time while including the interaction between depression risk group and brain activity. Q-values were estimated for target effects using the Benjamini-Hochberg method to adjust for false discovery rate (FDR).

After estimating the ANCOVA models, we conducted a series of integrative analyses to identify key biological pathways involved in relations between neural activity and gene expression. Effects were only retained for integrative analyses if (a) the q-value for the interaction between time and brain region was < 0.10 and (b) the log_2_-fold difference in gene expression change from baseline between one standard deviation above (high activity) and below (low activity) the mean of neural activity was ≥ 1, indicating at least a two-fold difference. We performed enrichment analyses of the final gene sets for each brain region/connection using the *pathfindR* R package and searched for enriched terms in the Kyoto Encyclopedia of Genes and Genomes (KEGG), Gene Ontology (GO), and Reactome databases. Pathways were only interpreted if the highest FDR-adjusted *p*-value across iterations was < 0.05. Hallmark immune gene sets from the Human Molecular Signatures Database (MSigDB) were also queried.

## 3. Results

### 3.1 Threat Network Activity and Gene Expression

#### 3.1.1 Summary of Primary Models

Results of the initial model that did not include the effects of individual differences in neural activity revealed statistically significant changes in expression of 1,528 genes from baseline to the post-task follow-ups. Across the target brain regions (ACC, amygdala, anterior insula, VMPFC) and their connections, we identified 18,924 statistically significant relationships between (*p*-value < 0.05) neural activity and change in gene expression, of which 10,775 met our 0.10 q-value and 2-fold change cut-offs (see Figure 2, panel A). Among the target brain regions (see Table 2), there were nearly twice as many transcripts related to activity of the anterior insula (*n* = 774) and VMPFC (*n* = 643) than the ACC (*n* = 442) and amygdala (*n* = 303). Furthermore, functional connectivity between the anterior insula and both the ACC (insula ACC: *n* = 1,572) and VMPFC (insula VMPFC: *n* = 1,587) were associated with differences in expression change of approximately 12% of the genes assessed, while other functional connection measures ranged from around 2-9%. Interestingly, the majority of transcripts related to insula-ACC connections were up-regulated when connectivity was high, while greater connectivity between the insula and VMPFC was associated with decreased expression of most genes, particularly at the second follow-up (see Figure 2, panel A).

**Fig. 2.**
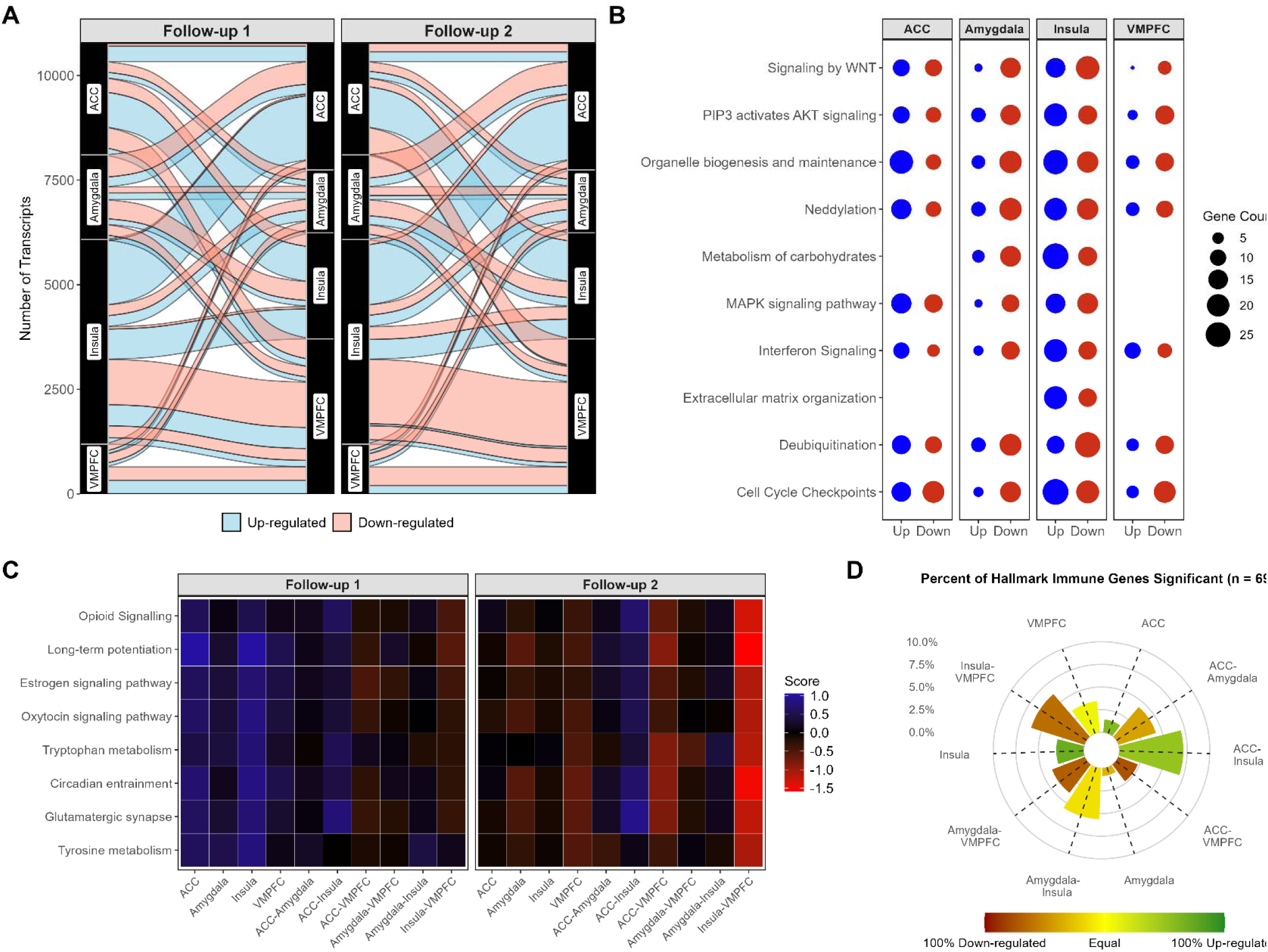
(A) Alluvial plot visualizing the number of significant effects at each time point for each brain region and connectivity between regions. Larger bars reflect a greater number of significant effects. Bars between the same region refer to univariate effects of region; bars connecting different regions refer to connectivity measures. (B) Dot plot showing the number of significant up- and own-regulated transcripts across general cell maintenance and defense pathways. (C) Agglomerated pathway enrichment scores (z-scored) accounting for the number of differentially up- and down-regulated genes in each neuroendocrine pathway, as well as the size of these effects. (D) Radial plot showing the percent of hallmark immune genes (MsigDB) differentially expressed for each brain region.

A similar pattern of results was found when surveying the hallmark immune gene set (*n* = 694 included in data; MsigDB). That is, insula-ACC and insula-VMPFC connectivity were related to 6-7% of canonical immune genes (Fig. 1, Panel D). Again, approximately 75% of the genes related to greater insula-ACC connectivity were up-regulated at either follow-up time point, while in contrast, the overwhelming majority of genes related to greater insula-VMPFC connectivity were down-regulated over time (81.40%). With the exception of functional connectivity between the anterior insula and amygdala, which was related to expression of 5.62% of immune genes, other regions and connections were associated with less than 5% of the gene set (see Figure 2, panel D).

#### 3.1.2 Gene Set Enrichment Analysis

Across the threat network regions and their connections, the majority of top enriched KEGG and Reactome pathway terms were related to general cell maintenance and communication (e.g., deubiquitination, MAPK signaling, cell cycle checkpoints), as is often found given the large available gene sets for these pathways. Although the major pathways enriched were similar, the number of differentially expressed genes and direction of relationships between neural activity and gene expression did vary across brain regions (see Figure 2, panel B). Analysis of agglomerated scores of top enriched terms related to neuroendocrine function (Figure 2, panel C) revealed that greater activity in the ACC and insula predicted up-regulation of genes related to long-term potentiation, oxytocin signaling, and tyrosine metabolism at the first follow-up, but these effects were absent or reversed at the second follow-up. Greater ACC-insula connectivity, on the other hand, was associated with up-regulation of pathways regulating dopaminergic and glutamatergic synapses at both follow-ups, among others. Moreover, greater ACC-VMPFC—and even more so insula-VMPFC—connectivity were related to down-regulation of most target neuroendocrine pathways, especially at follow-up 2.

#### 3.1.3 Target Immune Pathways

A survey of the top immune-specific enriched terms revealed several pathways related to threat network activity (see Fig. 3). Namely, greater activity in multiple brain regions, as well as greater connectivity between regions, were related to enrichment of key inflammatory pathways (e.g., hypoxia inducible factor-1 [HIF-1], IL-1, TNF), antiviral immunity (e.g., interferon signaling), and lymphocyte receptor signaling. Moreover, the degree of enrichment and proportion of up- vs. down-regulated genes in each pathway varied considerably by brain region. For example, with the ACC, anterior insula, and especially ACC-insula connectivity, greater activity was generally related to a predominance of up-regulated genes across immune pathways. In particular, over 85% of significant genes in the pro-inflammatory TNFR2 non-canonical NF- kB (fold enrichment: 2.19, *p* = 1.62 x 10^-15^), HIF-1 (fold enrichment: 1.17, *p* = 0.001), and interleukin-1 signaling pathways (fold enrichment: 2.50, *p* = 1.62 x 10^-16^) were up-regulated among those with high (compared to low) ACC-insula connectivity. Visualization of expression of individual genes in target immune pathways (Figure 3, panel B) further revealed especially large effect sizes for associations between higher ACC-insula activity and up-regulated genes in the interferon signaling pathway.

**Fig 3.**
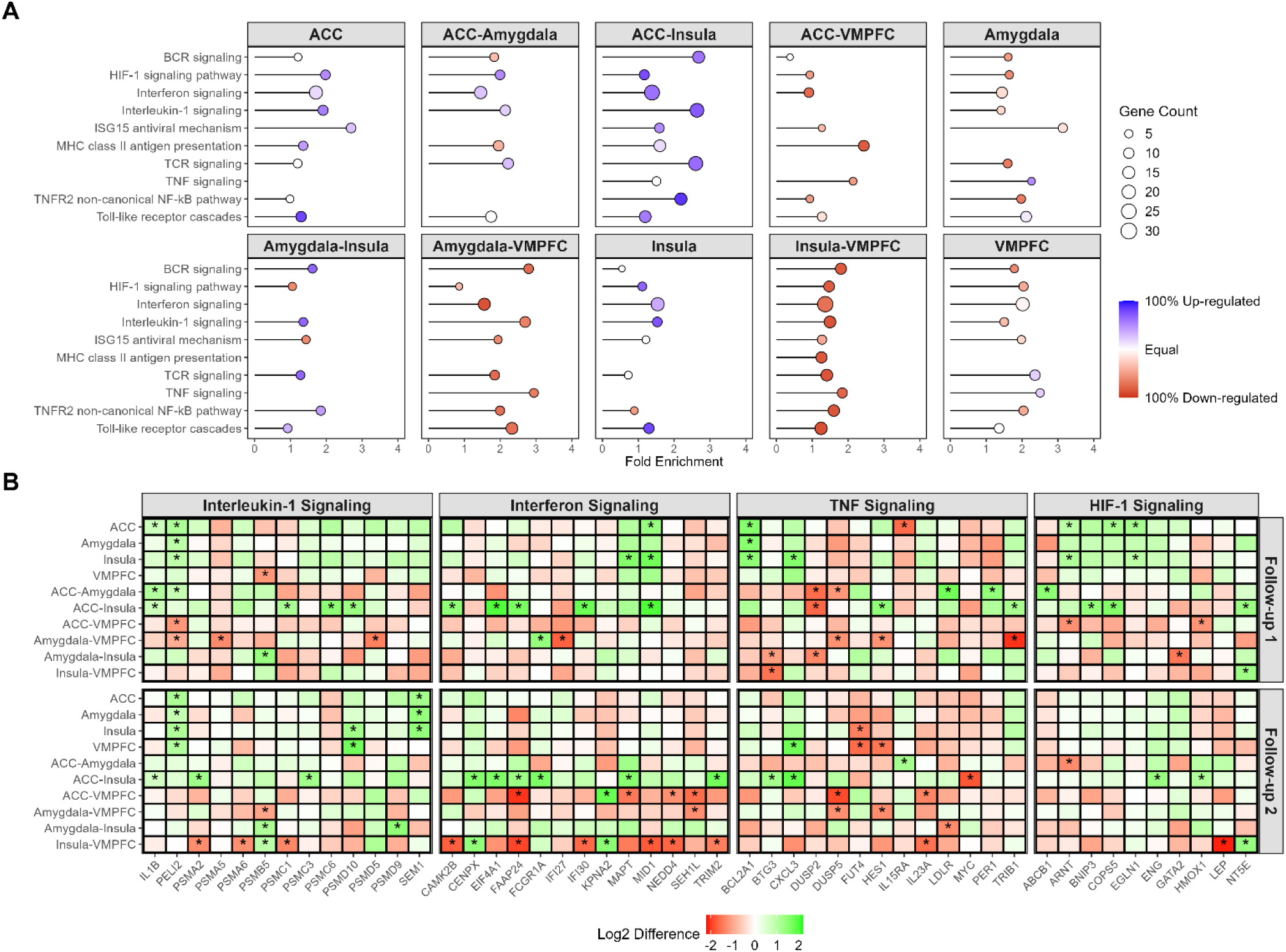
(A) Dot plot displaying the number of differently expressed genes for each brain region and key immune pathway, as well as the proportion of significant genes up- or down-regulated. (B) Heatmap showing log_2_-fold differences in expression of immune genes in key pathways between participants one standard deviation above and below the mean of each brain activity or connectivity measure.

While greater ACC activity was also associated with up-regulation of most key immune pathways, the highest proportion of up-regulated genes was found for toll-like receptor cascades (90%; fold enrichment: 1.30 *p* = 0.0009). On the other hand, in addition to up-regulation of interleukin-1 signaling (87.5% up-regulated; fold enrichment: 1.53, *p* = 0.00004) and toll-like receptor cascades (90% up-regulated; fold enrichment: 1.30, *p* = 0.0009) among other immune pathways, high insula activity predicted down-regulation of the TNFR2 NF-kB pathway, albeit weakly (75% down-regulated; fold enrichment: 0.89, *p* = 0.009). Interestingly, activities in both the ACC and insula were strongly associated with increased expression of multiple genes involved in HIF-1 signaling at follow-up 1 (e.g., *ARNT, BNIP3, EGLN1, COPS5*), but these effects diminished by the second follow-up. This pattern may suggest that elevated sensitivity to social threat is associated with transient enrichment of facets of the HIF-1 pathway under stress, which is consistent with its importance for the body’s response to myriad stressful conditions (e.g., hypoxia, oxidative stress, heat).^52–54^ Activity in the amygdala was associated with up-regulation of TNF signaling (75% up-regulated; fold enrichment: 2.26, *p* = 0.009), but down-regulation of other pathways, such as the TNFR2 NF-kB pathway (83.3% up-regulated; fold enrichment: 1.98, *p* = 0.00003) and HIF-1 signaling (80% up-regulated; fold enrichment: 1.65, *p* = 0.0006). On the whole, however, there were far fewer hits for amygdala activity alone than the amygdala’s connectivity to other regions.

Most notably, higher amygdala-VMPFC connectivity was linked to a majority of down-regulated genes for nearly all enriched immune pathways identified (see Figure S1), especially TNF (83.3% down-regulated; fold enrichment: 2.94, *p* = 0.000004), interleukin-1 (81.8% down-regulated; fold enrichment: 2.69, *p* = 9.36 x 10^-9^), and toll-like receptor signaling (85.7% down-regulated; fold enrichment: 2.33, *p* = 0.00001). Similar patterns of results were found for ACC-VMPFC and insula-VMPFC connectivity, with the latter offering perhaps the strongest case for domain-general down-regulation of immune pathways. In the latter case, we found no enriched immune pathways associated with insula-VMPFC connectivity that had a preponderance of up-regulated genes. Moreover, as is shown in Fig. 3, Panel B, the decreased expression of IL-1 and interferon signaling genes for participants with high (compared to low) insula-VMPFC connectivity was especially pronounced at follow-up 2. Interestingly, individuals with high insula-VMPFC connectivity (one standard deviation above the mean) demonstrated 4.5x greater decrease in expression of the gene encoding leptin than those with low connectivity (*LEP*; log_2_-fold difference: -2.125, *p* = 0.0009), which is involved in HIF-1 signaling in addition to its important role in energy regulation. Additionally, high insula-VMPFC connectivity was also associated with substantial decreases in *IL-23A* expression (log_2_-fold difference: -1.55, *p* = 0.005), as well as genes encoding complement proteins like *C1S* and *C2* (log_2_-fold differences: - 2.09 to -1.99, *p*s < 0.002).

#### 3.1.4 Interactions between Neural Activity and Depression Risk

Across brain activity and connectivity measures, we observed 13,430 statistically significant interactions with depression risk for at least one follow-up (log_2_-fold difference > 1 and *q* < 0.10, see Table S1). Among the univariate measures of brain activity, the majority of these interactions were found with the activities of the amygdala (*n* = 1391) and VMPFC (*n* = 1589). Enrichment analysis for genes associated with interactions between depression risk and brain activity is shown in Fig. 4, Panel A. Results revealed that for the VMPFC, genes for which the effects of brain activity differed by depression risk most often involved expression increasing alongside higher brain activity in both risk groups. This suggests that for most immune pathways, differences in VMPFC-gene expression relations between the high and low depression risk groups were found in the degree of up-regulation associated with greater VMPFC activity, not the direction of expression (i.e., up- vs. down-regulation). However, while the fold enrichment of TNF signaling associated with higher VMPFC activity was similar across risk groups, this pathway appeared to be primarily down-regulated among those with low depression risk (fold enrichment: 2.48, *p* = 0.003), and up-regulated for those with high risk (fold enrichment: 2.16, *p* = 0.002). Similarly, greater amygdala activity was related to up-regulated IL-1 signaling in the high risk group (fold enrichment: 1.41, *p* = 0.019), but down-regulated signaling in the low risk group (fold enrichment: 1.09, *p* = 0.002). Differences by depression risk group for other brain activity measures varied considerably by pathway.

**Fig. 4.**
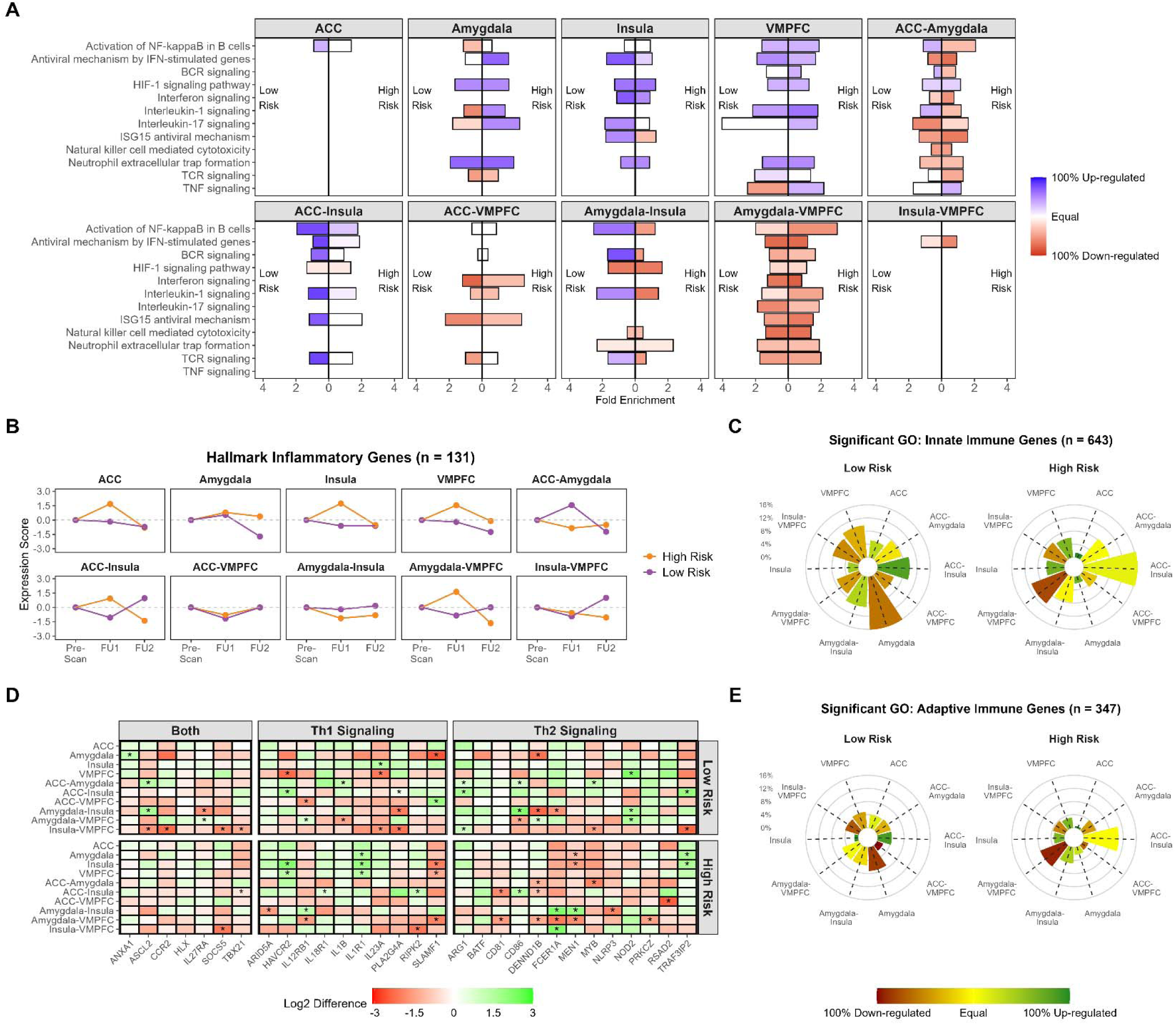
(A) Displays fold enrichment and proportion of genes up- or down-regulated in target immune pathways for individuals with high (bars moving right) and low depression risk (bars moving left). Results are shown for enrichment analysis of only genes with significant depression risk group by brain region interactions and thus reflect only genes for which the effects of brain activity on expression differ by risk group. (B) Plot visualizes a weighted mean of hallmark inflammatory genes (MsigDB) across follow-up time points for each brain region. Scores were calculated for each brain region by computing the mean of log_2_-fold difference values separately for genes up- and down-regulated, multiplying by the proportion of significant genes for each region up- or down-regulated and the proportion of total genes up- or down-regulated (*n* = 131), and then summing the scores for genes up- and down-regulated. (C) Shows heatmap of log_2_-fold differences for top differentially expressed genes in T helper cell (Th) 1 and 2 pathways by depression risk group. “Both” refers to genes included in both gene sets. (D) and (E) display radial plots indicating the percent of differentially expressed genes in the Gene Ontology (GO): Biological Process innate and adaptive immune pathways for each brain region.

With connectivity between brain regions, as was observed for the primary analyses, pathways impacted by significant interactions between risk group and amygdala-VMPC connectivity—and to a lesser extent ACC- and insula-VMPFC connectivity—nearly all involved a preponderance of down-regulated genes for both groups. That is, the effects of amygdala-VMPFC connectivity on immune gene expression differed by depression risk in the number or degree of down-regulated genes, not whether genes in a given pathway were primarily up- or down-regulated. For example, although higher amygdala-VMPFC connectivity was related to a greater fold enrichment of NF-kB activity in B cells for those with high depression risk (fold enrichment: 2.98, *p* = 0.000001) compared to low risk (fold enrichment: 1.99, *p* = 0.00003), in both cases, most significant genes in these pathways were down-regulated.

Furthermore, as is shown in Fig. 4, Panel B—which depicts changes in standardized expression scores for canonical inflammatory genes over time—enrichment for inflammatory signaling more broadly also varied by depression risk, brain activity, and time. Specifically, greater ACC, insula, and VMPFC activity, as well as ACC-insula and amygdala-VMPFC connectivity, were primarily associated with elevations in inflammatory gene expression at the first follow-up that returned to baseline (or slightly lower) approximately 30 minutes later at the second follow-up. In contrast, modest decreases in inflammatory gene expression were generally observed for the low risk group. The opposite pattern was found in relation to increased ACC-amygdala activity, whereby those with low depression risk exhibited elevated inflammatory gene expression at the first follow-up

Although the effects of interactions between brain activity and depression risk were relatively similar for T-helper cell 1 (Th1) and Th2 gene sets (Fig. 4, Panel C), a number of differences were observed when separately comparing genes related to innate and adaptive immune signaling (Fig 4., Panels D-E). Overall, brain activity was associated with differential expression of a greater proportion of innate than adaptive immune genes, regardless of depression risk. However, while amygdala activity was by far related to expression of the greatest percentage of innate immune genes among the low risk group (16.13% of gene set), most of which were down-regulated (80.36%), higher amygdala activity was only related to expression of 2% of innate immune genes in the high risk group (85.71% up-regulated). Conversely, greater ACC-insula connectivity was associated with expression of 16.42% of innate immune genes in the high risk group (56.14% up-regulated), but only 9.80% among participants with low depression risk (91.12% up-regulated). Moreover, consistent with previous findings, higher amygdala-VMPFC connectivity was related to differential expression of 10.95% of innate immune genes within the high risk group, the overwhelming majority of which were down-regulated (89.47%). For the most part, variation in adaptive immune gene expression across brain regions and depression risk resembled that found for innate immune genes, albeit with a smaller overall proportion of significant hits.

## 4. Discussion

The current research sought to examine whether individual differences in neural activity within the threat network, as well as functional connectivity between threat-related brain regions, impacted patterns of gene expression following a social evaluation task among adolescent females. Results revealed considerable variation in enrichment of diverse biological pathways associated with threat network neural activity and connectivity, providing some of the first evidence linking neural activity to transcriptional responses to social stress.

We found that consistent with previous research, elevated threat network activity was related to enrichment of several immune pathways, particularly those associated with innate immunity and inflammation.^27, 49^ In particular, greater functional connectivity between the ACC and anterior insula was generally related to up-regulation of most immune pathways. Prior research suggests that these regions are involved in representing visceral states and processing peripheral inflammatory signals,^55–58^ which may help explain their consistent activation during emotionally salient and socially evaluative experiences,^26, 49, 59^ as well as their role in stress-related autonomic and immune responses. In contrast, greater connectivity between the insula and VMPFC was broadly associated with down-regulation, especially for interferon, IL-1, and HIF-1 signaling. A similar pattern emerged for amygdala–VMPFC connectivity, which was also linked to broad downregulation across nearly all major immune pathways. The VMPFC has been implicated in processes related to executive control and emotion regulation, ^60–62^ and its engagement during social evaluation has been interpreted as a potential mechanism for attenuating affective and physiological responses. In light of existing research and our findings, this pattern may suggest that VMPFC coupling contributes to broader dampening of stress-related physiological reactivity, including transcriptional immune responses. It is important to emphasize, however, any inferences about underlying psychological processes remain speculative and should be viewed as plausible, but not definitive.

While activation in individual brain regions influenced immune gene expression, these effects appeared to vary by signaling pathway, suggesting a context-dependent or function-specific role in shaping immune responses to social threat. Notably, activity in both the ACC and insula was associated with elevated expression of several genes involved in HIF-1 signaling (e.g., *ARNT, BNIP3, EGLN1, COPS5*) at the first follow-up, though this association waned by the second. This transient upregulation may reflect an acute, time-sensitive response to social threat, consistent with HIF-1’s established role in regulating cellular adaptation to stressors such as hypoxia, oxidative stress, and inflammation.^52–54^

The present study also aimed to examine whether depression risk moderated relationships between neural activity and gene expression. Indeed, we found that depression risk affected which brain regions and connectivity measures were primarily related to gene expression, as well as the timing, degree, and even direction of transcriptional changes within key immune pathways. For example, among participants with low depression risk, greater amygdala activity was associated with down-regulation of a large portion of innate immune genes, while the opposite was true for the high-risk group — a pattern that may reflect altered threat calibration among youth with increased vulnerability to depression. This is an interesting extension on previous research which has shown that high-risk adolescents who exhibit greater amygdala activation in response to social evaluation report more pronounced increases in depressed mood and social disconnection than their low-risk peers.^45^ Moreover, while the direction of transcriptional effects linked to connectivity patterns was largely consistent across risk groups, notable differences emerged in their magnitude. Specifically, ACC–VMPFC, Amygdala– VMPFC, and Insula–VMPFC connectivity were associated with more pronounced downregulation of immune gene expression in the high-risk group. Further investigation into magnitude-based differences may clarify whether they represent a heightened biological sensitivity to social threat in high-risk adolescents—an idea central to the Social Signal Transduction Theory of Depression.^11^

Although immune pathways were the primary focus, additional analyses revealed transcriptional changes in broader cellular and neuroendocrine pathways. Top enriched KEGG and Reactome terms included deubiquitination, MAPK signaling, and cell cycle regulation— common findings in transcriptome-wide studies. We also found region- and connectivity-specific effects on neuroendocrine pathways (e.g., oxytocin, dopamine, glutamate), particularly at follow-up 1 for ACC and insula activity, and more sustained for ACC–insula and insula–VMPFC connectivity. While secondary to our aims, these findings may inform future research on depression-related comorbidities.

Several limitations warrant consideration. Although the sample size was sufficient to detect moderate effects and replicate prior findings, it may have lacked power to identify smaller or more complex patterns, and some effect sizes could be inflated. The demographically diverse sample was drawn from a WEIRD population, potentially limiting generalizability. While maternal depression history was confirmed through structured interviews, undetected subclinical symptoms in the low-risk group cannot be fully excluded. Participation in a neuroimaging and social evaluation study may have also introduced stress-related variability unrelated to depression risk. Lastly, gene expression results were not validated at the protein level, and future studies should examine whether these transcriptional changes yield downstream functional effects.

Despite these limitations, the present findings offer valuable insight into the neural and transcriptional correlates of social threat processing in adolescence. Taken together, the findings contribute to a growing body of research examining how neural responses to social threat may relate to biological processes implicated in depression. By examining both neural activity/connectivity and functional genomic responses in the same individuals, this study helps to address key gaps identified in the Social Signal Transduction Theory of Depression.^11^ Future research will be important for clarifying the temporal dynamics and clinical relevance of these associations, as well as their potential role in the emergence of depressive symptoms over time.

## Supplemental Tables

**Table 1.**
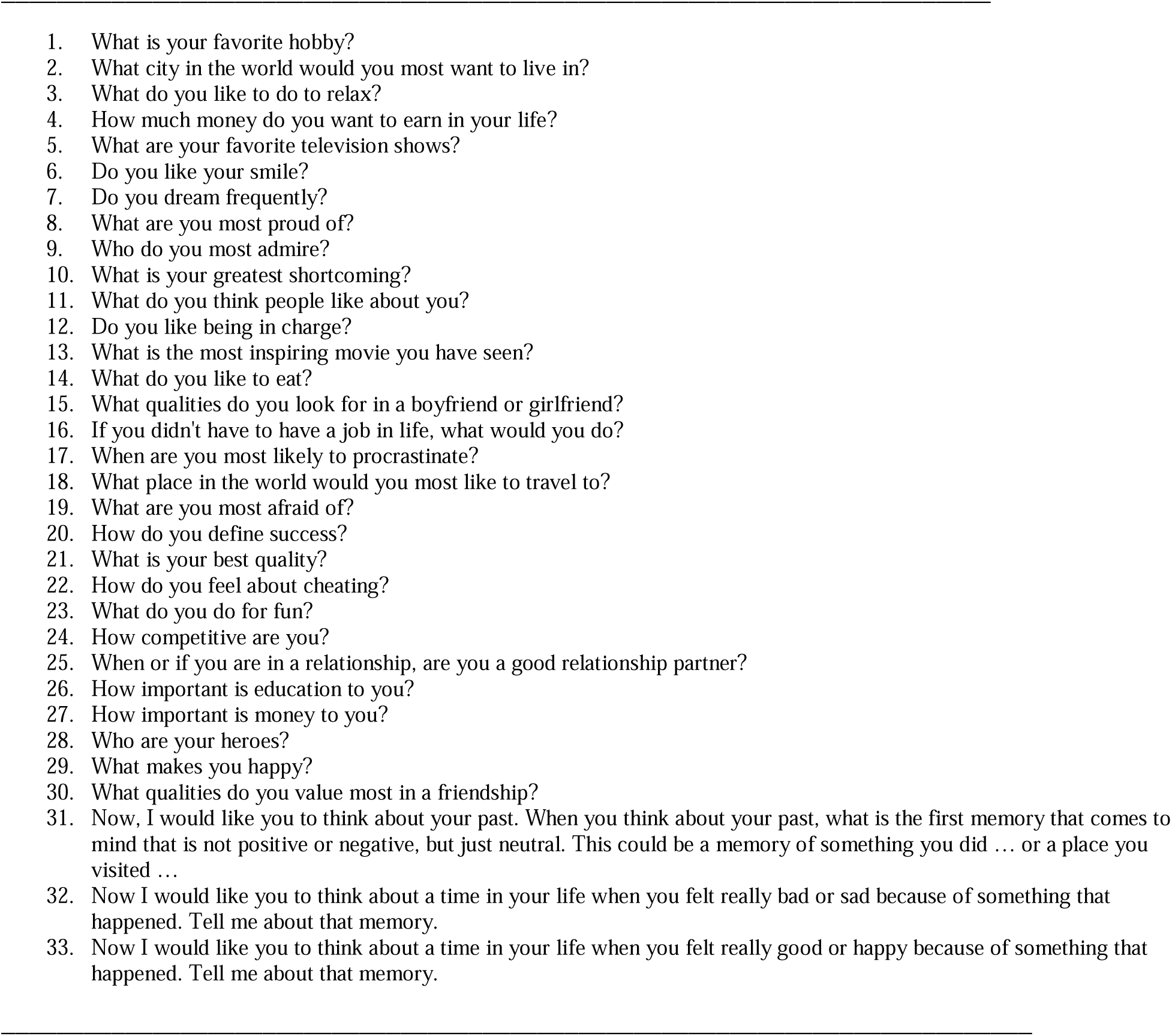
Interview questions assessing daughters’ interests, opinions, values, & childhood memories.

## References

1. Meyer, I. H. Prejudice, social stress, and mental health in lesbian, gay, and bisexual populations: Conceptual issues and research evidence. Psychol. Bull. 129, 674–697 (2003).

2. Sichko, S. et al. Psychobiology of Stress and Adolescent Depression (PSY SAD) Study: Protocol overview for an fMRI-based multi-method investigation. Brain Behav. Immun. - Health 17, 100334 (2021).

3. Slavich, G. M., Thornton, T., Torres, L. D., Monroe, S. M. & Gotlib, I. H. Targeted Rejection Predicts Hastened Onset of Major Depression. J. Soc. Clin. Psychol. 28, 223–243 (2009).

4. Monroe, S. M., Rohde, P., Seeley, J. R. & Lewinsohn, P. M. Life events and depression in adolescence: Relationship loss as a prospective risk factor for first onset of major depressive disorder. J. Abnorm. Psychol. 108, 606–614 (1999).

5. Slavich, G.M., 2016. Psychopathology and Stress. in The SAGE Encyclopedia of Theory in Psychology (SAGE Publications, Inc., 2455 Teller Road, Thousand Oaks, California 91320, 2016). doi:10.4135/9781483346274.n262.

6. Slavich, G. M., O’Donovan, A., Epel, E. S. & Kemeny, M. E. Black sheep get the blues: A psychobiological model of social rejection and depression. Neurosci. Biobehav. Rev. 35, 39– 45 (2010).

7. Nolan, S. A., Flynn, C. & Garber, J. Prospective relations between rejection and depression in young adolescents. J. Pers. Soc. Psychol. 85, 745–755 (2003).

8. Mojtabai, R., Olfson, M. & Han, B. National Trends in the Prevalence and Treatment of Depression in Adolescents and Young Adults. Pediatrics 138, e20161878 (2016).

9. Salk, R. H., Hyde, J. S. & Abramson, L. Y. Gender differences in depression in representative national samples: Meta-analyses of diagnoses and symptoms. Psychol. Bull. 143, 783–822 (2017).

10. Slavich, G. M. & Cole, S. W. The Emerging Field of Human Social Genomics. Clin. Psychol. Sci. 1, 331–348 (2013).

11. Slavich, G. M. & Irwin, M. R. From stress to inflammation and major depressive disorder: A social signal transduction theory of depression. Psychol. Bull. 140, 774–815 (2014).

12. Furman, D. et al. Chronic inflammation in the etiology of disease across the life span. Nat. Med. 25, 1822–1832 (2019).

13. Slavich, G. M., Mondelli, V. & Moriarity, D. P. Psychoneuroimmunology of depression. in APA handbook of depression: Vol. 1. Classification, co-occurring conditions, and etiological processes vol. 1 (American Psychological Association, in press).

14. Slavich, G. M. Psychoneuroimmunology of Stress and Mental Health. in The Oxford Handbook of Stress and Mental Health (eds. Harkness, K. L. & Hayden, E. P.) 518–546 (Oxford University Press, 2020). doi:10.1093/oxfordhb/9780190681777.013.24.

15. Slavich, G. M. Social Safety Theory: A Biologically Based Evolutionary Perspective on Life Stress, Health, and Behavior. Annu. Rev. Clin. Psychol. 16, 265–295 (2020).

16. Slavich, G. M. & Sacher, J. Stress, sex hormones, inflammation, and major depressive disorder: Extending Social Signal Transduction Theory of Depression to account for sex differences in mood disorders. Psychopharmacology (Berl.) 236, 3063–3079 (2019).

17. Boyce, W. T., Sokolowski, M. B. & Robinson, G. E. Toward a new biology of social adversity. Proc. Natl. Acad. Sci. 109, 17143–17148 (2012).

18. McEwen, B. S. & Gianaros, P. J. Central role of the brain in stress and adaptation: Links to socioeconomic status, health, and disease. Ann. N. Y. Acad. Sci. 1186, 190–222 (2010).

19. Nelson, E. E., Leibenluft, E., McCLURE, E. B. & Pine, D. S. The social re-orientation of adolescence: a neuroscience perspective on the process and its relation to psychopathology. Psychol. Med. 35, 163–174 (2005).

20. Ho, T. C. et al. Functional connectivity of negative emotional processing in adolescent depression. J. Affect. Disord. 155, 65–74 (2014).

21. Kaiser, R. H., Andrews-Hanna, J. R., Wager, T. D. & Pizzagalli, D. A. Large-Scale Network Dysfunction in Major Depressive Disorder: A Meta-analysis of Resting-State Functional Connectivity. JAMA Psychiatry 72, 603 (2015).

22. Dedovic, K., Slavich, G. M., Muscatell, K. A., Irwin, M. R. & Eisenberger, N. I. Dorsal Anterior Cingulate Cortex Responses to Repeated Social Evaluative Feedback in Young Women with and without a History of Depression. Front. Behav. Neurosci. 10, (2016).

23. Hamilton, J. P. et al. Functional Neuroimaging of Major Depressive Disorder: A Meta-Analysis and New Integration of Baseline Activation and Neural Response Data. Am. J. Psychiatry 169, 693–703 (2012).

24. Gong, L. et al. Disrupted reward circuits is associated with cognitive deficits and depression severity in major depressive disorder. J. Psychiatr. Res. 84, 9–17 (2017).

25. Silk, J. S. et al. Subgenual Anterior Cingulate Cortex Reactivity to Rejection Vs. Acceptance Predicts Depressive Symptoms among Adolescents with an Anxiety History. J. Clin. Child Adolesc. Psychol. 52, 659–674 (2023).

26. Eisenberger, N. I., Taylor, S. E., Gable, S. L., Hilmert, C. J. & Lieberman, M. D. Neural pathways link social support to attenuated neuroendocrine stress responses. NeuroImage 35, 1601–1612 (2007).

27. Slavich, G. M., Way, B. M., Eisenberger, N. I. & Taylor, S. E. Neural sensitivity to social rejection is associated with inflammatory responses to social stress. Proc. Natl. Acad. Sci. 107, 14817–14822 (2010).

28. Muscatell, K. A., Merritt, C. C., Cohen, J. R., Chang, L. & Lindquist, K. A. The Stressed Brain: Neural Underpinnings of Social Stress Processing in Humans. in Neuroscience of Social Stress (eds. Miczek, K. A. & Sinha, R.) vol. 54 373–392 (Springer International Publishing, Cham, 2021).

29. Berretz, G., Packheiser, J., Kumsta, R., Wolf, O. T. & Ocklenburg, S. The brain under stress—A systematic review and activation likelihood estimation meta-analysis of changes in BOLD signal associated with acute stress exposure. Neurosci. Biobehav. Rev. 124, 89–99 (2021).

30. Vijayakumar, N., Cheng, T. W. & Pfeifer, J. H. Neural correlates of social exclusion across ages: A coordinate-based meta-analysis of functional MRI studies. NeuroImage 153, 359– 368 (2017).

31. Wang, H., Braun, C. & Enck, P. How the brain reacts to social stress (exclusion) – A scoping review. Neurosci. Biobehav. Rev. 80, 80–88 (2017).

32. Slavich, G. M., Mengelkoch, S. & Cole, S. W. Human social genomics: Concepts, mechanisms, and implications for health. Lifestyle Med. 4, e75 (2023).

33. Leschak, C. J. & Eisenberger, N. I. Two Distinct Immune Pathways Linking Social Relationships With Health: Inflammatory and Antiviral Processes. Psychosom. Med. 81, 711–719 (2019).

34. Miller, G. E., Chen, E. & Parker, K. J. Psychological stress in childhood and susceptibility to the chronic diseases of aging: Moving toward a model of behavioral and biological mechanisms. Psychol. Bull. 137, 959–997 (2011).

35. Dieckmann, L., Cole, S. & Kumsta, R. Stress genomics revisited: gene co-expression analysis identifies molecular signatures associated with childhood adversity. Transl. Psychiatry 10, 34 (2020).

36. Slavich, G. M. Social Safety Theory: Understanding social stress, disease risk, resilience, and behavior during the COVID-19 pandemic and beyond. Curr. Opin. Psychol. 45, 101299 (2022).

37. Slavich, G. M. et al. Social Safety Theory: Conceptual foundation, underlying mechanisms, and future directions. Health Psychol. Rev. 17, 5–59 (2023).

38. Cain, D. W. & Cidlowski, J. A. Immune regulation by glucocorticoids. Nat. Rev. Immunol. 17, 233–247 (2017).

39. Kenney, M. J. & Ganta, C. K. Autonomic Nervous System and Immune System Interactions. in Comprehensive Physiology (ed. Terjung, R.) 1177–1200 (Wiley, 2014). doi:10.1002/cphy.c130051.

40. Mengelkoch, S., Alley, J. C., Cole, S. W. & Slavich, G. M. Transcriptional evidence of HPA axis dysregulation in adolescent females: Unique contributions of chronic early-life stressor exposure and maternal depression history. J. Affect. Disord. 371, 245–252 (2025).

41. Mengelkoch, S. & Slavich, G. M. Sex Differences in Stress Susceptibility as a Key Mechanism Underlying Depression Risk. Curr. Psychiatry Rep. 26, 157–165 (2024).

42. Irwin, M. R. et al. Moderators for depressed mood and systemic and transcriptional inflammatory responses: a randomized controlled trial of endotoxin. Neuropsychopharmacology 44, 635–641 (2019).

43. Quinn, M. E., Stanton, C. H., Slavich, G. M. & Joormann, J. Executive Control, Cytokine Reactivity to Social Stress, and Depressive Symptoms: Testing the Social Signal Transduction Theory of Depression. Stress 23, 60–68 (2020).

44. Seiler, A., Von Känel, R. & Slavich, G. M. The Psychobiology of Bereavement and Health: A Conceptual Review From the Perspective of Social Signal Transduction Theory of Depression. Front. Psychiatry 11, 565239 (2020).

45. Shields, G. S. et al. Heightened neural activity and functional connectivity responses to social rejection in female adolescents at risk for depression: Testing the Social Signal Transduction Theory of Depression. J. Affect. Disord. 345, 467–476 (2024).

46. Slavich, G. M. et al. Interpersonal life stress, inflammation, and depression in adolescence: Testing Social Signal Transduction Theory of Depression. Depress. Anxiety 37, 179–193 (2020).

47. O’Connor, M.-F. et al. To assess, to control, to exclude: Effects of biobehavioral factors on circulating inflammatory markers. Brain. Behav. Immun. 23, 887–897 (2009).

48. Lieb, R., Isensee, B., Höfler, M., Pfister, H. & Wittchen, H.-U. Parental Major Depression and the Risk of Depression and Other Mental Disorders in Offspring: A Prospective-Longitudinal Community Study. Arch. Gen. Psychiatry 59, 365 (2002).

49. Muscatell, K. A. et al. Greater amygdala activity and dorsomedial prefrontal–amygdala coupling are associated with enhanced inflammatory responses to stress. Brain. Behav. Immun. 43, 46–53 (2015).

50. Eisenberger, N. I., Inagaki, T. K., Muscatell, K. A., Byrne Haltom, K. E. & Leary, M. R. The Neural Sociometer: Brain Mechanisms Underlying State Self-esteem. J. Cogn. Neurosci. 23, 3448–3455 (2011).

51. Bhanji, J., Smith, D. V. & Delgado, M. A Brief Anatomical Sketch of Human Ventromedial Prefrontal Cortex. Preprint at 10.31234/osf.io/zdt7f (2019).

52. Chen, S. & Sang, N. Hypoxia-Inducible Factor-1: A Critical Player in the Survival Strategy of Stressed Cells. J. Cell. Biochem. 117, 267–278 (2016).

53. Ely, B. R., Lovering, A. T., Horowitz, M. & Minson, C. T. Heat acclimation and cross tolerance to hypoxia: Bridging the gap between cellular and systemic responses. Temperature 1, 107–114 (2014).

54. Lee, P., Chandel, N. S. & Simon, M. C. Cellular adaptation to hypoxia through hypoxia inducible factors and beyond. Nat. Rev. Mol. Cell Biol. 21, 268–283 (2020).

55. Capuron, L. et al. Anterior Cingulate Activation and Error Processing During Interferon-Alpha Treatment. Biol. Psychiatry 58, 190–196 (2005).

56. Craig, A. D. How do you feel? Interoception: the sense of the physiological condition of the body. Nat. Rev. Neurosci. 3, 655–666 (2002).

57. Eisenberger, N. I., Inagaki, T. K., Rameson, L. T., Mashal, N. M. & Irwin, M. R. An fMRI study of cytokine-induced depressed mood and social pain: The role of sex differences. NeuroImage 47, 881–890 (2009).

58. Harrison, N. A. et al. Neural origins of human sickness in interoceptive responses to inflammation. Biol. Psychiatry 66, 415–422 (2009).

59. Wager, T. D. et al. Brain mediators of cardiovascular responses to social threat, Part II: Prefrontal-subcortical pathways and relationship with anxiety. NeuroImage 47, 836–851 (2009).

60. Kerestes, R., Davey, C. G., Stephanou, K., Whittle, S. & Harrison, B. J. Functional brain imaging studies of youth depression: A systematic review. NeuroImage Clin. 4, 209–231 (2014).

61. Lamm, C. & Lewis, M. D. Developmental Change in the Neurophysiological Correlates of Self-Regulation in High- and Low-Emotion Conditions. Dev. Neuropsychol. 35, 156–176 (2010).

62. Wagner, D. D. & Heatherton, T. F. Self-regulatory depletion increases emotional reactivity in the amygdala. Soc. Cogn. Affect. Neurosci. 8, 410–417 (2013).

